# Pan-cancer machine learning predictors of primary site of origin and molecular subtype

**DOI:** 10.1101/333914

**Authors:** William F. Flynn, Sandeep Namburi, Carolyn A. Paisie, Honey V. Reddi, Sheng Li, R. Krishna Murthy Karuturi, Joshy George

## Abstract

**Background:** It is estimated by the American Cancer Society that approximately 5% of all metastatic tumors have no defined primary site (tissue) of origin and are classified as *<u>c</u>ancers of <u>u</u>nknown <u>p</u>rimary* (CUPs). The current standard of care for CUP patients depends on immunohistochemistry (IHC) based approaches to identify the primary site. The addition of post-mortem evaluation to IHC based tests helps to reveal the identity of the primary site for only 25% of the CUPs, emphasizing the acute need for better methods of determination of the site of origin. CUP patients are therefore given generic chemotherapeutic agents resulting in poor prognosis. When the tissue of origin is known, patients can be given site specific therapy with significant improvement in clinical outcome. Similarly, identifying the primary site of origin of metastatic cancer is of great importance for designing treatment.

Identification of the primary site of origin is an import first step but may not be sufficient information for optimal treatment of the patient. Recent studies, primarily from The Cancer Genome Atlas (TCGA) project, and others, have revealed molecular subtypes in several cancer types with distinct clinical outcome. The molecular subtype captures the fundamental mechanisms driving the cancer and provides information that is essential for the optimal treatment of a cancer. Thus, along with primary site of origin, molecular subtype of a tumor is emerging as a criterion for personalized medicine and patient entry into clinical trials.

However, there is no comprehensive toolset available for precise identification of tissue of origin or molecular subtype for precision medicine and translational research.

**Methods and Findings:** We posited that metastatic tumors will harbor the gene expression profiles of the primary site of origin of the cancer. Therefore, we decided to learn the molecular characteristics of the primary tumors using the large number of cancer genome profiles available from the TCGA project. Our predictors were trained for 33 cancer types and for the 11 cancers where there are established molecular subtypes. We estimated the accuracy of several machine learning models using cross-validation methods. The extensive testing using independent test sets revealed that the predictors had a median sensitivity and specificity of 97.2% and 99.9% respectively without losing classification of any tumor. Subtype classifiers achieved median sensitivity of 87.7% and specificity of 94.5% via cross validation and presented median sensitivity of 79.6% and specificity of 94.6% in two external datasets of 1,999 total samples. Importantly, these external data shows that our classifiers can robustly predict the primary site of origin from external microarray data, metastatic cancer data, and patient-derived xenograft (PDX) data.

**Conclusion:** We have demonstrated the utility of gene expression profiles to solve the important clinical challenge of identifying the primary site of origin and the molecular subtype of cancers based on machine learning algorithms. We show, for the first time to our knowledge, that our pan-cancer classifiers can predict multiple cancers’ primary site of origin from metastatic samples. The predictors will be made available as open source software, freely available for academic non-commercial use.

## 1. INTRODUCTION

Precision cancer therapy requires the knowledge of primary site of origin and accurate subtyping of the cancer to identify an appropriate therapeutic regimen. However, according to the American Cancer Society, an estimated 2 to 5 percent of all cancer patients have metastatic tumors for which routine testing cannot locate the primary site and is therefore classified as a cancer of unknown primary (CUP). CUP patients have very poor prognosis, primarily because the course of treatment is empiric and not tailored for a specific tumor type [1, 2]. In addition, lack of knowledge of the true cancer type puts CUP patients under severe psychological distress that may lead to clinically significant depressive symptoms [3].

In a study of CUP patients that were predicted to have primary tumors originating in the colon, the median survival of patients increased in those that received site-specific chemotherapy as compared to those who received empirically-determined treatments [4, 5]. Furthermore, multiple studies have supported the use of molecular profiling to diagnose CUP and to determine specific treatment based upon the predicted site of origin leading to an improvement in overall survival [6–9]. Therefore, it is important to develop systematic methods to identify the primary site of origin of the disease.

Currently, immunohistochemistry (IHC) utilizing antibodies targeted to certain tumor-specific antigens is the main method for primary site identification in patients with CUP [2, 10]. However, there is not a single specific marker that can be used to conclusively diagnose the primary tumor, leading to the use of multiple different IHC markers, and generating the possibility that different clinicians will arrive at different diagnoses of the primary tumor type [2, 11–14]. Recent advances in genomics has led to the development of potential new means for diagnosing these tumors; multiple studies have utilized gene expression profiles and other molecular markers to diagnose CUP [2]. One such method involved comparisons of gene expression profiles between CUPs and a set of primary and metastatic tumors with known origins to predict the tissue of origin for the CUP [2, 6, 15–20]. None of these tools were able to identify the tissue of origin with high sensitivity and specificity for all the CUP samples. For example, the EPICUP tool, which had the best performance to date provides high accuracy for only 87% of the CUP samples.

Several studies, including the Cancer Genome Atlas (TCGA) and International Cancer Genome Consortium (ICGC) studies, have shown that the cancers from the primary tissue can be classified into molecular subtypes with distinct clinical outcome and therapeutic options [21–28]. The molecular subtype information can also have predictive power. For example, Bevacizumab, a monoclonal antibody that block angiogenesis, is shown to benefit patients of mesenchymal and proliferative subtypes in ovarian cancer and therefore may be used as a criterion for the entry into a clinical trial [29]. In addition, molecular subtype information can guide the selection of targeted therapies and to suggest new treatment strategies (e.g. the potential use of JAK2 inhibitors and PD-L1/2 antagonists for the treatment of EBV-positive gastric cancer) [27]. However, identification of molecular subtypes is clinically challenging and thus clinicians are unable to utilize molecular subtype information to inform treatment decisions [30, 31] due to a lack of tools and assays for pan-cancer subtyping despite the availability of genomic technologies for clinical diagnostics.

To fill this important gap in the clinical and translational research setting, using expression data available from the TCGA project, we developed Machine Learning based predictors that enable accurate identification of primary site of origin and subtype of cancer (Figure 1). Our predictors were trained for 33 cancer types and for the 11 cancers where there are established molecular subtypes. The extensive testing using independent test sets revealed that the predictors had a median sensitivity and specificity of 97.2% and 99.9% respectively without failing to classify any tumor using in total 1,959 samples. Subtype predictors achieved median sensitivity of 79.6% and specificity of 94.6% using two external validation sets consisting of 1,999 breast and ovarian cancer samples in total. Our gene expression-based pan-cancer classifier can, for the first time to our knowledge, robustly predict multiple cancers’ primary site of origin from metastatic samples using an independent validation dataset (specificity: 99.3%, sensitivity: 82.1%) and predict molecular subtype. Compared to other pan-cancer classifiers based on somatic mutations, our classifier is not limited to only cancer types with high mutation burden and has much greater potential for clinical diagnosis and therapeutic design.

**Figure 1.**
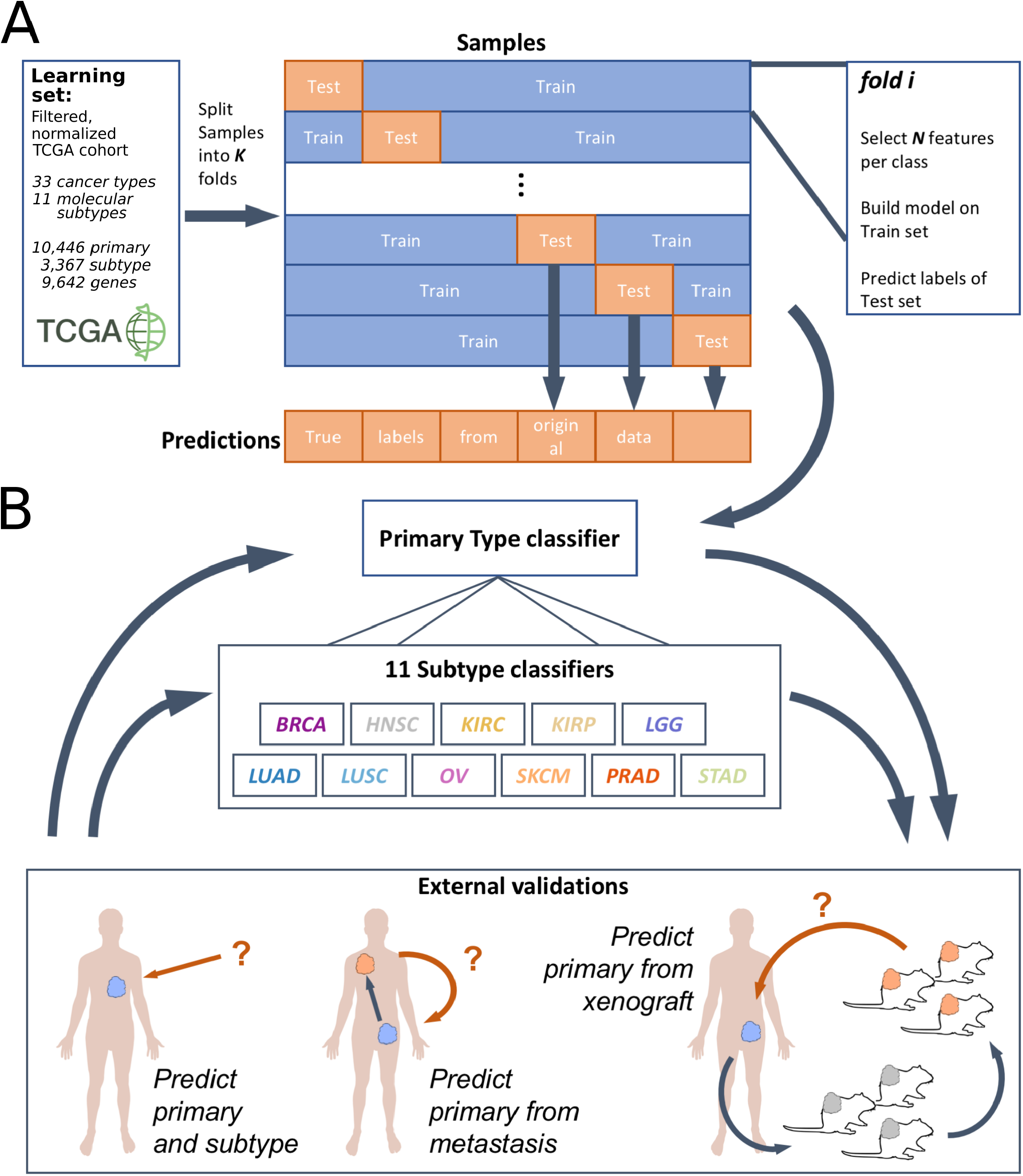
Platform-independent learning and validation from TCGA transcriptomes. (A) Schematic showing the learning procedure used to train machine learning models from labeled TCGA transcriptomes spanning 33 cancer types and 11 molecular subtypes. Models were trained and evaluated using k-fold cross validation on normalized and standard scaled expression profiles. For each fold, N features were selected from each class (see Methods) and pooled, which were used to train the classification model. Using this schema, we constructed one primary type and eleven molecular subtype predictors for each type of model: random forest (RF), support vector machine (SVM), k-nearest neighbor classifier (kNN), and diagonal linear discriminant analysis (DLDA). (B) Classification performance was evaluated via cross-validation on the learning set and external validation utilizing five datasets; using two of these datasets, we challenged the predictors to infer primary tumor types from transcriptomes of metastatic or passaged patient-derived xenograft samples.

## 2. METHODS

### 2.1 Expression datasets

#### 2.1.1 Learning set: TCGA expression data

RSEM [32] normalized mRNA expression matrices were downloaded for each of the 33 unique cancer cohorts (listed in Table 1) available from the Broad Institute GDAC Firehose (run 2016_01_28) [33]. Individual cohort expression matrices were converted to Biobase ExpressionSet objects [34] for standardization and then combined as an ExpressionSet stored in Apache Feather format (version 0.4.0) to allow downstream analysis in both R and Python (https://github.com/wesm/feather). The raw expression matrix consisted 11,330 samples and 20,531 genes, which was reduced to 10,446 samples and 9,642 genes after filtering (detailed below).

**Table 1.** 33 GDAC Cancer Cohorts

| Cohort Abbr. | Cases | Disease Name |
| --- | --- | --- |
| ACC | 92 | Adrenocortical carcinoma |
| BLCA | 412 | Bladder urothelial carcinoma |
| BRCA | 1,098 | Breast invasive carcinoma |
| CESC | 307 | Cervical and endocervical cancers |
| CHOL | 51 | Cholangiocarcinoma |
| COAD | 460 | Colon adenocarcinoma |
| DLBC | 58 | Lymphoid Neoplasm Diffuse Large B-cell Lymphoma |
| ESCA | 185 | Esophageal carcinoma |
| GBM | 613 | Glioblastoma multiforme |
| HNSC | 528 | Head and Neck squamous cell carcinoma |
| KICH | 113 | Kidney Chromophobe |
| KIRC | 537 | Kidney renal clear cell carcinoma |
| KIRP | 323 | Kidney renal papillary cell carcinoma |
| LAML | 200 | Acute Myeloid Leukemia |
| LGG | 516 | Brain Lower Grade Glioma |
| LIHC | 377 | Liver hepatocellular carcinoma |
| LUAD | 585 | Lung adenocarcinoma |
| LUSC | 504 | Lung squamous cell carcinoma |
| MESO | 87 | Mesothelioma |
| OV | 602 | Ovarian serous cystadenocarcinoma |
| PAAD | 185 | Pancreatic adenocarcinoma |
| PCPG | 179 | Pheochromocytoma and Paraganglioma |
| PRAD | 499 | Prostate adenocarcinoma |
| READ | 171 | Rectum adenocarcinoma |
| SARC | 261 | Sarcoma |
| SKCM | 470 | Skin Cutaneous Melanoma |
| STAD | 443 | Stomach adenocarcinoma |
| TGCT | 150 | Testicular Germ Cell Tumors |
| THCA | 503 | Thyroid carcinoma |
| THYM | 124 | Thymoma |
| UCEC | 560 | Uterine Corpus Endometrial Carcinoma |
| UCS | 57 | Uterine Carcinosarcoma |
| UVM | 80 | Uveal Melanoma |

#### 2.1.2 External validation data

External cancer gene expression datasets, i.e. those used exclusively for testing the performance of models, were obtained from the publicly available Gene Expression Omnibus (GEO) and the patient-derived xenograft (PDX) mouse models generated at the Jackson Laboratory. These datasets were not introduced during model fitting (hence external validation) and were generated using both RNA-seq and microarray technologies.

The first dataset used to test the model accuracy was the microarray gene expression profile of 2,158 cancer samples from the expression project for oncology (expO, GSE2109) [35]. Of the 2,158 samples, we were able to identify relevant primary tumor types for 1,558 samples and excluded LGG/GBM from classification due to too few samples, resulting in classification of 1,552 samples. The second dataset, GSE18549, contained expression profiles of 96 tumors from their metastatic sites [36]. We used 88 of these tumors whose primary sites could be identified and those with more than 1 sample per primary site. The third dataset used contains the expression profile of 338 PDX RNA-seq samples generated at the Jackson Laboratory and available through the Mouse Tumor Biology (MTB) gene expression portal (http://tumor.informatics.jax.org) [37]. Of these 338 samples, 325 samples could be mapped to one of the 33 TCGA primary cancer types and the 7 OV samples were excluded as they originate from the same two patients. The distribution of the primary types in the external datasets used for validation are shown in Table 2.

**Table 2.** Distribution of samples in the three external datasets used to validate the primary classification model

| Primary site | External dataset |  |  |
| --- | --- | --- | --- |
|  | GSE2109 | GSE18549 | PDX |
| BLCA | 32 | 0 | 29 |
| BRCA | 354 | 14 | 41 |
| COAD | 312 | 35 | 68 |
| DLBC | 0 | 0 | 4 |
| KIRC/KIRP/KIRH | 281 | 6 | 8 |
| LAML | 0 | 0 | 13 |
| LGG/GBM | 6 | 0 | 5 |
| LIHC | 46 | 0 | 0 |
| LUAD/LUSC | 133 | 10 | 88 |
| OV | 279 | 14 | 7 |
| PAAD | 0 | 0 | 12 |
| PRAD | 83 | 9 | 0 |
| READ | 0 | 1 | 0 |
| SARC | 0 | 0 | 32 |
| SKCM | 0 | 0 | 18 |
| THCA | 32 | 0 | 0 |
| N/A | 600 | 7 | 13 |
| **Used** | 1,552 | 88 | 318 |
| **Total** | 2,158 | 96 | 338 |

For external validation of our subtype predictors, we acquired two additional microarray datasets. The first, accession number GSE9899, contains 215 ovarian cancer samples [15] and the second, EGA study EGAS00000000083 (https://www.ebi.ac.uk/ega), contains 1,784 breast cancer samples [38]. Both datasets comprise 4 molecular subtypes each: mesenchymal, immunoreactive, differentiated, and proliferative for the ovarian set; and basal-like, HER2-enriched, luminal A, and luminal B for the breast set.

#### 2.1.3 Normalization, filtering, and preprocessing

All expression data was log2-transformed, and only genes with (a) maximum log2 expression greater than 8 and (b) variance in log2 expression greater than 1 were retained. After filtering, the genes in each dataset was scaled to zero mean expression and unit variance. This scaling allows expression to be measured in terms of standard deviations and affords platform-independent use of subsequently trained models.

#### 2.1.4 Molecular subtype label curation

Molecular subtype information was downloaded from cBioPortal [39, 40] for 3,367 samples from the following primary cancers: glioblastoma multiforme (GBM), stomach adenocarcinoma (STAD), breast (BRCA), ovarian (OV), prostate (PRAD), and lung squamous cell cancers (LUSC). Further annotations were curated from the following supplemental data files: lower grade glioma (LGG) [41], head and neck squamous cell carcinoma (HNSC) [22], uterine corpus endometrial carcinoma (UCEC) [42], cutaneous melanoma (SKCM) [43], papillary (KIRP) [44] and clear cell (KIRC) [45] renal cancers, and lung adenocarcinoma (LUAD) [27]. R (version 3+) scripts were written to extract relevant information (e.g. sample id, specific subtype) from downloaded data and supplemental files. These scripts are available in the public project GitHub repository as described under Code availability.

#### 2.1.5 Pan-organ group labels

A recently published study performed integrated clustering on the multiomic data from the approximately 10,000 The Cancer Genome Atlas (TCGA) samples of 33 types of cancer. The authors identified multi-cancer groups, which tend to span whole organs or related organ groups [46]. We use these pan-organ group assignments to evaluate our classification results in a individual- or multi-organ context. These classifications are reproduced here: central nervous system (GBM LGG), core gastrointestinal (ESCA, STAD, COAD, READ), developmental gastrointestinal (LIHC, PAAD, CHOL), endocrine (THCA and ACC), gynecologic (OV, UCEC CESC BRCA), head and neck (HNSC), hematologic and lymphatic malignancies (LAML, DLBC, THYM), melanocytic (SKCM and UVM), neural-crest-derived tissues (PCPG), soft tissue (SARC and UCS), thoracic (LUAD, LUSC, MESO), urologic (BLCA, PRAD, TGCT, KIRC, KICH, KIRP).

### 2.2 Machine Learning Algorithms for Cancer Classification

We evaluated several popular machine learning algorithms to develop predictors for CUP classification and subtype identification: DLDA, KNN, SVM and Random Forest. We used R-packages *sparsediscrim* (version 0.2.4)*, base::knn, e1071* (version 1.6-8), and *randomForest* (version 4.6-14), respectively for the training, testing; and *caret* (version 6.0-79) for tool development. Unless otherwise specified, default parameters were chosen for model construction.

#### 2.2.1 Diagonal Linear Discriminant Analysis (DLDA)

The DLDA classifier belongs to the family of Naive Bayes classifiers, where the distribution of each class is assumed to be a multivariate normal and to share a common covariance matrix. The DLDA classifier is a modification to LDA, where the off-diagonal elements of the pooled sample covariance matrix are set to zero [47]. DLDA was used as classifier in several genomic based cancer classification tasks [48, 49].

#### 2.2.2 k-Nearest Neighbor (KNN) classifier

A KNN classifier offers the simplest classifier training strategy, also referred to as ‘lazy classifier’, and has been successfully applied in the classification of cancer and non-cancer related classification tasks [50–52]. A KNN classifier uses the training samples as reference vectors and, for every sample in the test set, the k nearest (in Euclidean distance) reference vectors are found. The classification is decided by majority vote of the k-nearest neighbors’ class. Note that, if multiple nearest neighbor vectors are found with identical distances, all such nearest neighbor vectors are included in the voting pool, which can lead to k being exceeded in these cases [53].

#### 2.2.3 Support Vector Machine (SVM)

An SVM algorithm builds a predictive model by constructing a representation of the training samples as points in higher dimensional space and builds a linear model (separating linear boundary) in that space such that the mapped samples of the different categories are separated by a gap that is as wide as possible. New examples are then mapped into that same space and predicted to belong to a category based on which side of the separating boundary they fall. For multiclass classification among N classes, N*(N-1)/2 binary SVM classifiers are constructed and trained in a one-versus-one manner; ultimate class predictions come from voting amongst the ensemble of binary classifiers. The implementation used for our classifiers employed a Gaussian kernel. SVMs were successfully used for classification of samples in variety of studies [54–56].

#### 2.2.4 Random Forest

The random forest algorithm employs a collection of decision trees constructed from bootstrapped input data and classification is done by majority voting among the ensemble of trees [57]. Single decision trees are prone to overfitting; multiple trees constructed from randomly sampled copies of the input data allows the consensus classification to be robust and extensible to new samples. Each of our random forest models constructed 1000 trees, each tree constructed from randomly sampled input with replacement, and each decision tree node uses 31 randomly selected features to partition the tree.

### 2.3 Model training and external validation

The schema for predictor design for primary site classification and subtype classification are depicted in Figure 1.

#### 2.3.1 Design of primary site (tumor type) predictor

All models were trained using the same feature selection and cross-validation schedule. Each model was then trained using a 3-fold cross validation procedure as follows. The expression set is partitioned into 3 random subsamples and for each partition: (1) the selected partition is used as the testing set and the remaining 2 are combined into a training set; (2) the 100 most differentially expressed genes in each class (cancer type) are selected, measured by log-fold-change of differential expression between in-class and out-of-class samples (p < 0.001); (3) the model is trained using the selected features; (4) predictions for the selected partition is recorded. The cross-validation procedure yields an estimate of the model performance with the selected parameters. The final model is then constructed using the entire set of samples and 1,971 unique genes selected via the procedure in (2) above.

#### 2.3.2 Molecular subtype classification

For each of the 11 primary cancer types with established molecular subtypes, a model is constructed as described above using the scaled, log2-transformed expression of the sample corresponding to the selected primary type as input. For each cancer type, similar to cancer type classification, features are selected by computing the differential expression (p<0.001) in each subtype in comparison with the other subtypes of the same cancer type.

#### 2.3.3 Predictor performance metrics

Each classification algorithm (predictor) was compared using per class and overall positive predictive value, sensitivity, and specificity. Additional metrics such as per-class balance accuracy, and F1 score are included in the supplementary tables. Per class metrics are computed using a one-versus-all scheme.

### 2.4 Visualization

An interactive web application was constructed using the Python Dash framework (version 0.21.0). The application shows the TCGA data embedded in three dimensions using UMAP (umap-learn, version 0.2.1, [58]) and t-SNE (MulticoreTSNE, version 0.1, [59, 60]). Data points are color coded by tumor type or primary site (with cancer and match normal samples), and molecular subtype, which can be controlled interactively through the interface.

The web application and the associated code are freely available for non-commercial, academic use at https://pccportal.jax.org (pan-cancer classification portal) and the source code is available as described in the Code availability subsection.

### 2.5 Code availability

All code to download, process, train, and validate these data and models is available in the following GitHub repository: https://github.com/TheJacksonLaboratory/tcga_subtype_classification. All results and the figures can be easily reproduced by cloning and running make.

## 3. RESULTS

### 3.1 Precise classification of primary cancer types across platforms

The classification of the 33 primary cancer types from the TCGA cohort (9,642 samples) by random forest is presented in Figure 2A. Classification yields a median sensitivity and specificity of 97.2% and 99.9% (n=33), respectively (Figure 2C). The major misclassifications are primarily within organ systems (Figure 2B). Indeed, when primary types are grouped by pan-organ groups [46], the median sensitivity increases to 98.5% with a substantial improvement in the minimum sensitivity to 86.0% (Figure 2B,D). It is important to note that every sample was classified by our model; no samples were excluded from classification, either by a sample quality metric or through lack of consensus during label assignment. The most frequent misclassifications occur between nearby locations in the gastrointestinal tract: rectal adenocarcinoma (READ) is completely misclassified as colon adenocarcinoma (COAD), and esophageal carcinoma (ESCA) is often misclassified as stomach adenocarcinoma (STAD).

**Figure 2.**
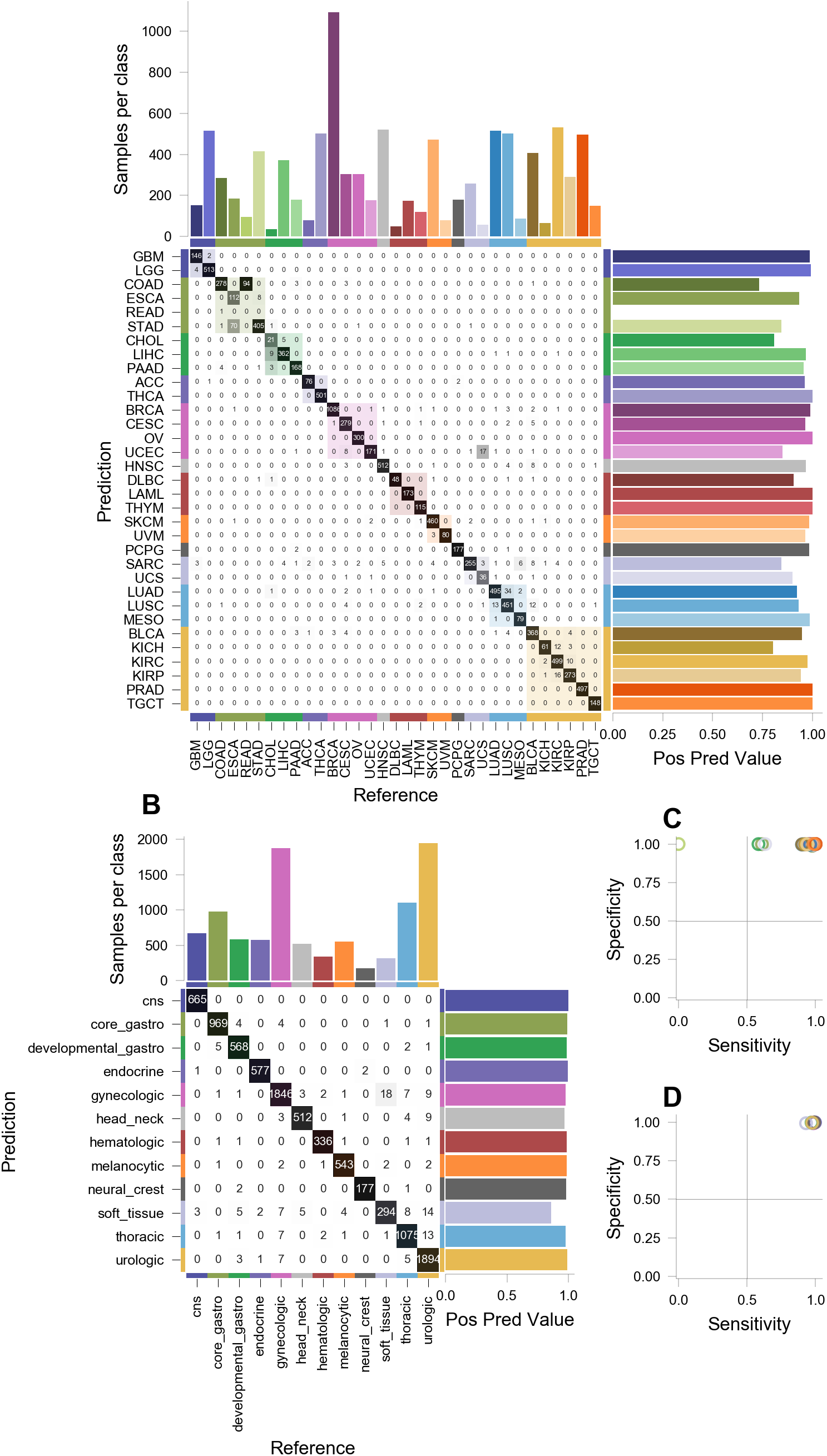
Precise classification of tumors by primary type and organ system. (A-B) Random forest classification of primary cancer types and grouped by pan-organ system. Text in contingency table cell *c_j,i_* shows tumor of class *i* classified as class *j*. Grayscale shading of table cells is proportional to the number of samples of each primary site, represented as bars above the table. Color shading along the main diagonal shows pan-organ groups. Positive predictive value (precision) for each prediction class are shown to the right of the table. (C-D) Sensitivity and specificity for each classification in (A) and (B), respectively.

COAD and READ are so similar that they are typically considered as a single primary type, colorectal carcinoma (CRC) [23]. Misclassification between ESCA and STAD is expected, as a certain class of ESCAs (esophageal adenocarcinomas) present at the interface of the esophagus and stomach [61]. Also of note is the misclassification of uterine carcinosarcoma (UCS) as uterine corpus endometrial carcinoma (UCEC). Histologically, USC presents features of both UCEC and sarcoma (SARC) [62].

To understand these misclassifications, the expression profiles of every training sample was embedded into a two-dimensional latent space using UMAP (see Methods) and colored by primary tumor type, shown in Figure 3. Several anatomical and histological structures readily emerge from the embedding. Some cancers are observed to form disparate, well-separated clusters by organ system, such as brain (GBM-LGG), liver and gallbladder (LIHC, CHOL), and kidneys (KIRC, KIRP, KIRH), while other cancers are grouped by histological features, such as the melanomas (SKCM, UVM) and squamous cell cancers (BLCA, CECS, HNSC, LUSC, and some ESCA) forming distinct clusters. The core gastrointestinal tract cancers cluster tightly, with COAD and READ embedded into an inseparable mass which is adjoined by STAD and some ESCA samples. ESCA samples clearly segregate into two populations, consistent with both esophageal adenocarcinoma (clustered with STAD) and squamous cell carcinoma (clustered with LUSC, HNSC, etc.) being classified under ESCA (Zheng 2013). Similarly, the known similarities between USC, UCEC, and SARC clearly emerges, with the embedding of USC forming a bridge between UCEC and SARC clusters; we also observe two distinct clusters of SARC samples, one most similar to USC and the other most similar to UCEC. As this embedding is heavily dependent on the input samples and number thereof, it may be that some misclassifications are unavoidable without a larger cohort of samples.

**Figure 3.**
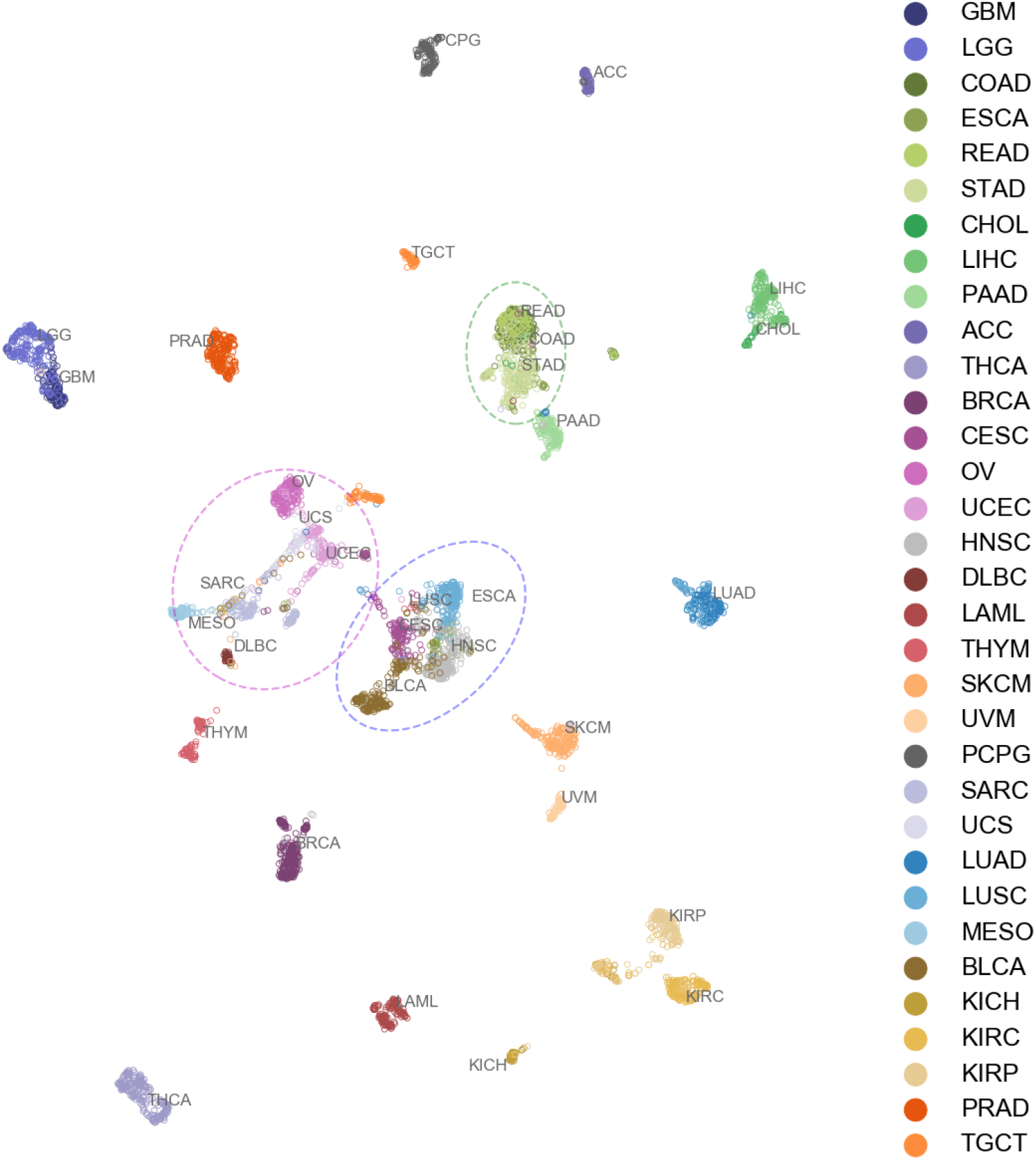
Unsupervised embedding of expression profiles reveals relationships among primary sites. Expression profiles from all samples were embedded into two dimensions using uniform manifold approximation and projection (UMAP) [58] and colored by primary cancer type. For each cancer, labels are placed near the centroid of the expression profile in the UMAP latent space. Anatomical and histological relationships are emergent and add context to the most common misclassifications in Figure 2. The following groups of cancers are highlighted with green, blue, and purple ellipses, respectively: COAD, READ, STAD; BLCA, CESC, ESCA, HNSC, LUSC; OV, SARC, UCEC, UCS.

#### 3.1.1 Primary site predictor performed well on external expression data

To further validate the primary site predictor, we classified 1,552 samples across 9 primary cancer types (Figure 4A) profiled using microarrays from the Expression Project for Oncology (expO, GSE2109 [35]). Due to the age of the dataset, primary cancer types corresponding to the brain (LGG, GBM), lung (LUAD, LUSC), and kidney (KIRC, KIRP, KIRH) were aggregated to match the respective primary site annotations of the dataset. Further, the genes used for classification were reduced to 1,788 genes from the initially selected genes of 1,971 in order to match those found in the external dataset, and the model was retrained on the training set with only these 1,788 features. This external validation set not only tests the predictor performance independent of batch variation, but also its independence of the platform and robustness to feature loss which are critical for the application of the predictors in clinical and translational research.

**Figure 4.**
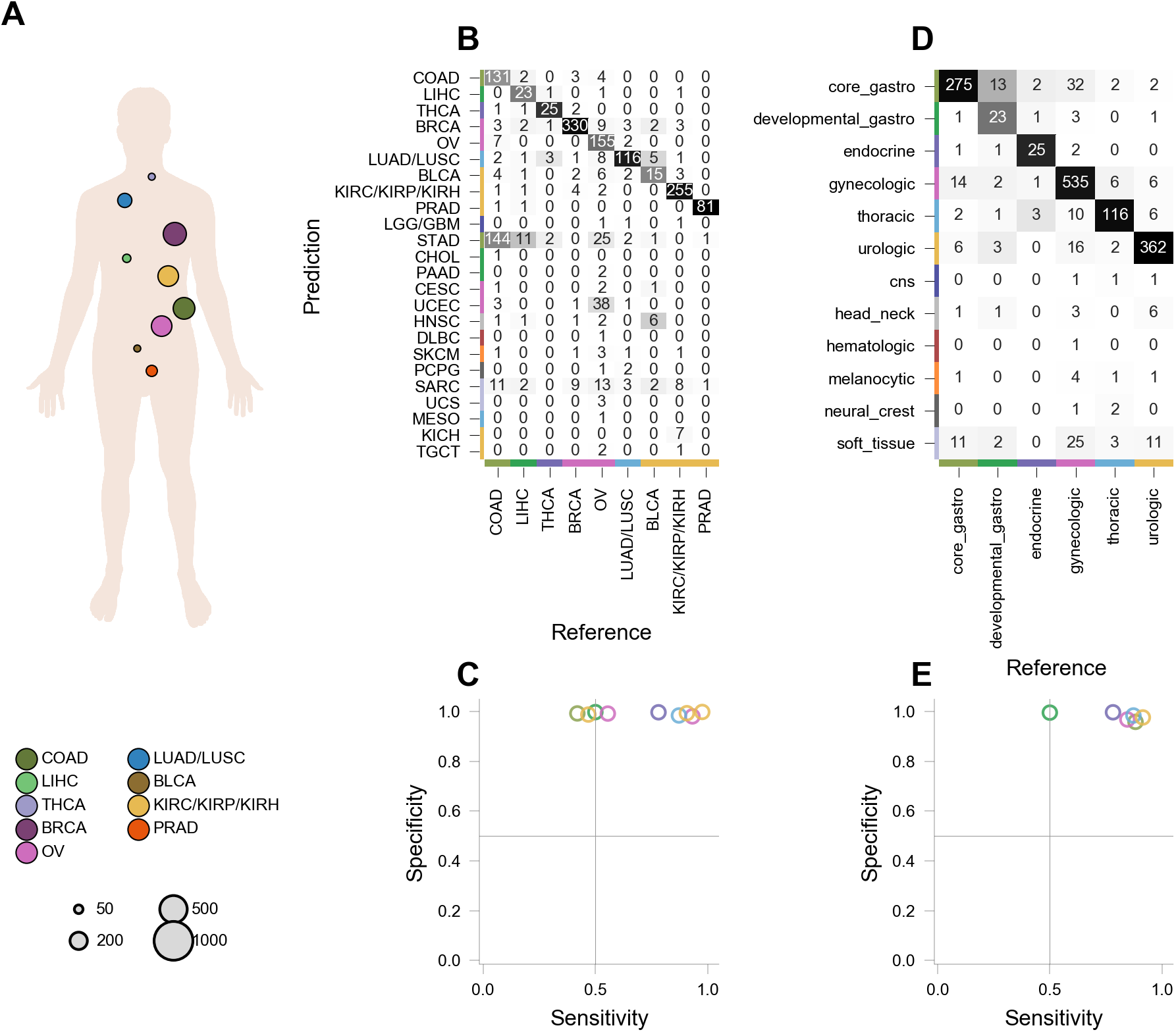
External validation of primary site predictor using microarray data. 1,552 microarray expression profiles (GSE2109) from 9 cancer types or related cancer types (LUSC/LUAD and KIRC/KIRP/KICH) were classified to further validate the primary site predictor (A). Markers in (A) are scaled by the number of samples in each class. Classification was evaluated by primary site (B) and pan-organ group (C). (D-E) Sensitivity and specificity for each classification in (B) and (C), respectively.

Classification by primary site, shown in Figure 4B, yields median specificity of 99.3% and median sensitivity of 78.1% (n=9) (Figure 4C), with misclassifications largely within organ systems. For example, misclassification arises between gastrointestinal cancers STAD, COAD, and LIHC, and ovarian serous cystadenocarcinoma (OV) is misclassified as cancers with similar histology or anatomical location. When the classification is reorganized by pan-organ group, as shown in Figure 4D, median sensitivity increases to 86.0% (n=6) with the misclassification only between core and developmental gastrointestinal cancers (Figure 4E).

#### 3.1.2 Primary site predictor can identify cancer of unknown primary

Identification of the primary cancer (site) of origin from a metastatic sample is a significant clinical challenge. As metastases are expected to retain the transcriptional signature of primary tumor of origin, we hypothesize that our predictors can identify the primary tumor type from metastatic samples. We examined the performance of our predictors using an external validating dataset from metastatic tumors.

Primary site classification of expression profiles of 88 metastatic samples across 6 known primary sites is shown in Figure 5A,B [36]. The median specificity is 99.3%, and the median sensitivity is 82.1%. The most common misclassifications were, again, between COAD and STAD, the vast majority involving metastatic tumors in the liver. Between-organ-system classification (Figure 5C) shows the minimum sensitivity substantially improves from 20.0% to 69.0%, where predictions of these gastrointestinal cancers are combined (Figure 5D). Further examination revealed that the most common metastases in the data, 30/88 (34%) samples, are to the liver or lung from the colon, illustrated in Figure 5E. Of the 88 tumors, 52 metastases (59.1%) are classified to the correct primary tumor type, 72 metastases (81.8%) are classified to the correct primary organ system, 7 metastases (7.9%) are classified as the tumor from the respective metastasized sites, and 9 metastases (10.1%) are classified incorrectly (Figure 5F) i.e. neither as tumor of primary site nor as the metastasized site.

**Figure 5.**
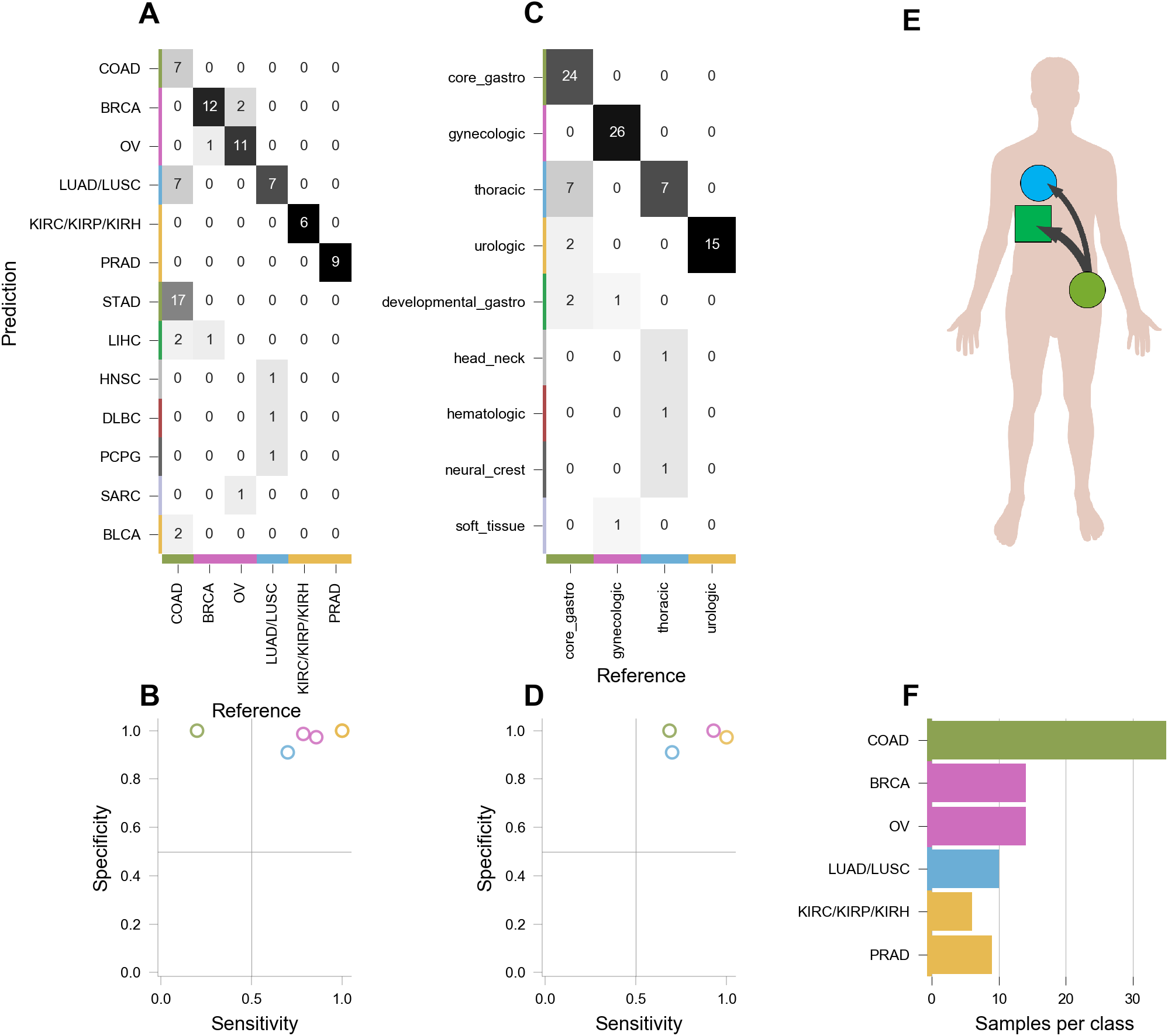
Predictor infers primary cancer of origin from metastatic tumor samples. (A-B) 88 expression profiles of metastatic tumors (GSE18549) from primary site of origin spanning 6 organs were classified by primary cancer type and primary pan-organ group. (C-D) Sensitivity and specificity for each classification in (A) and (B), respectively. (E) 30/88 (34%) of samples are liver or lung metastases from the colon. (F) The majority of misclassifications of primary site are within pan-organ system; of the remainder, 7 misclassifications identify the metastatic tumor whereas 9 are true misclassifications.

We further validate our predictors independently using an external dataset from patient-derived xenograft (PDX) models of cancer. PDX models of cancer are a great resource to evaluate therapeutic regimens but can also be used as a tool to study metastatic cancer, as illustrated in Figure 6A. Primary tumor is resected from human patients and tumor fragments are implanted into a cohort of immunodeficient mice [63]. After a growth period, the mouse-grown tumor is resected and implanted into a new generation of mice. This process can be repeated several times.

**Figure 6.**
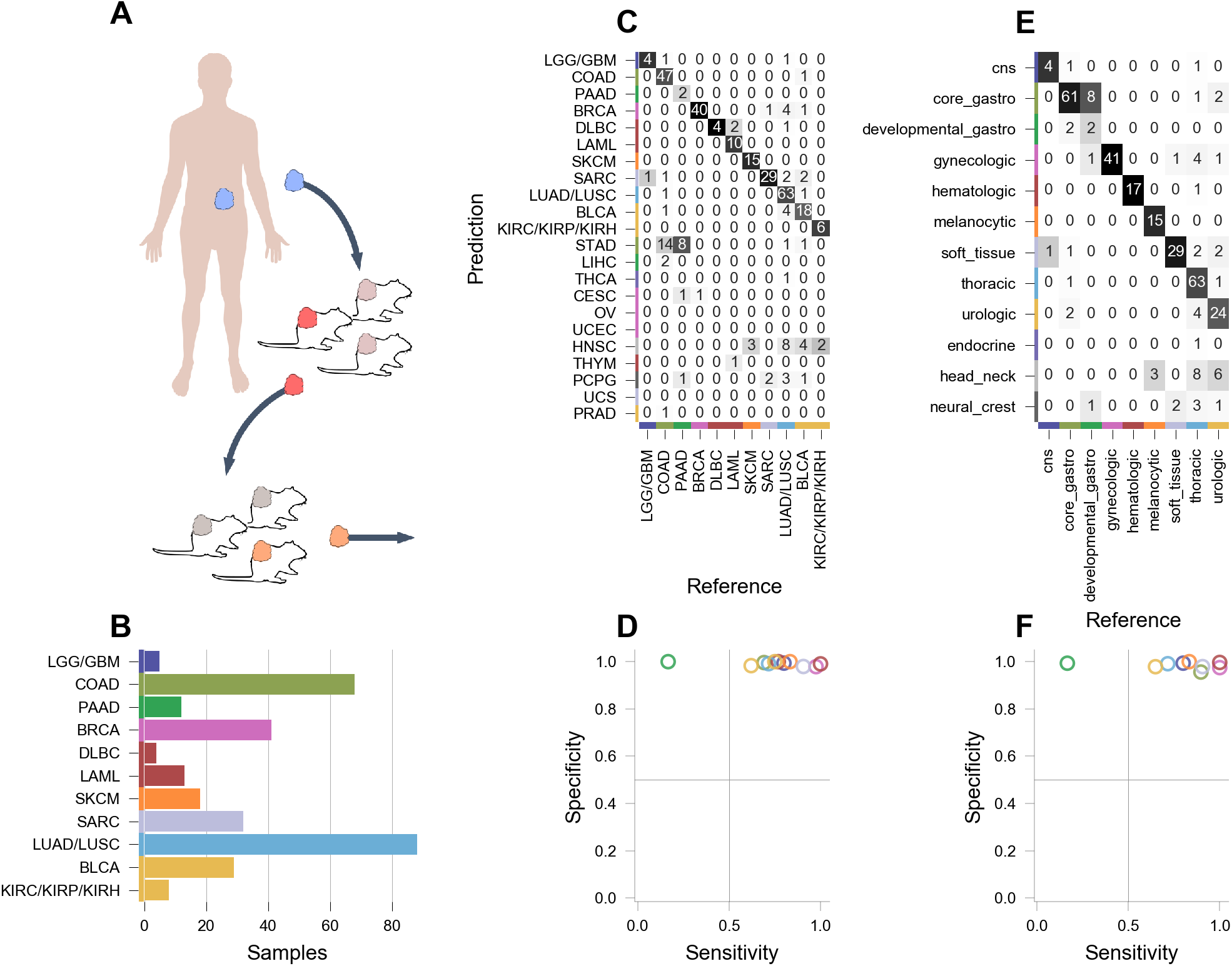
Predictor infers primary cancer of origin from passaged patient-derived xenografts. (A) 295 expression profiles of resected passaged patient-derived xenograft (PDX) tumors from primary sites spanning 11 organs were classified by primary site (samples generated at the Jackson Laboratory, available via MTB [37]). PDX tumor samples were taken for sequencing from the second generation of mice. Classification of primary site identification was evaluated by primary site (B) and pan-organ group (C). (D-E) Sensitivity and specificity for each classification in (B) and (C), respectively.

We performed the primary site type classification of 318 PDX-derived mouse-grown tumors (samples were taken from the second generation of mice) spanning 11 primary sites (Figure 6B-C). Classification of primary cancer types yields a median specificity of 99.5% (n=11), and median sensitivity of 76% (n=11) (Figure 6D). When classified by pan-organ system, the median sensitivity increases to 83% (Figure 6E,F). Despite not being present in the set of primary cancers, several tumors including COAD and PAAD are classified as STAD, possibly due to the close proximity of anatomic positions.

These three external validations of our model overwhelmingly support the hypothesis that metastatic and xenograft tumors retain the molecular signature of the primary tumor.

Application of such models in the clinic will allow for more effective treatments of CUPs.

### 3.2 Subtype specific classification accurately identifies molecular subtypes

Molecular subtypes have been defined for 11 cancer types: BRCA, HNSC, KIRC, KIRP, LGG, LUAD, LUSC, OV, PRAD, SKCM, and STAD. Each of these primary types has two to four molecular subtypes. For example, breast cancers are frequently subtyped into Basal-like, Her2-enriched, Luminal A and Luminal B. This subtyping has prognostic power and can be used as predictive marker for therapeutic approaches [64].

Eleven models were constructed, one model for each primary tumor type, into its molecular subtypes, as illustrated schematically in Figure 1. The positive predictive value, sensitivity, specificity, and number of samples per subtype are shown in Figure 7A-D. The best performing subtype predictors, LGG, LUAD, PRAD, have median sensitivity above 90%, with PRAD yielding nearly perfect classification.

**Figure 7.**
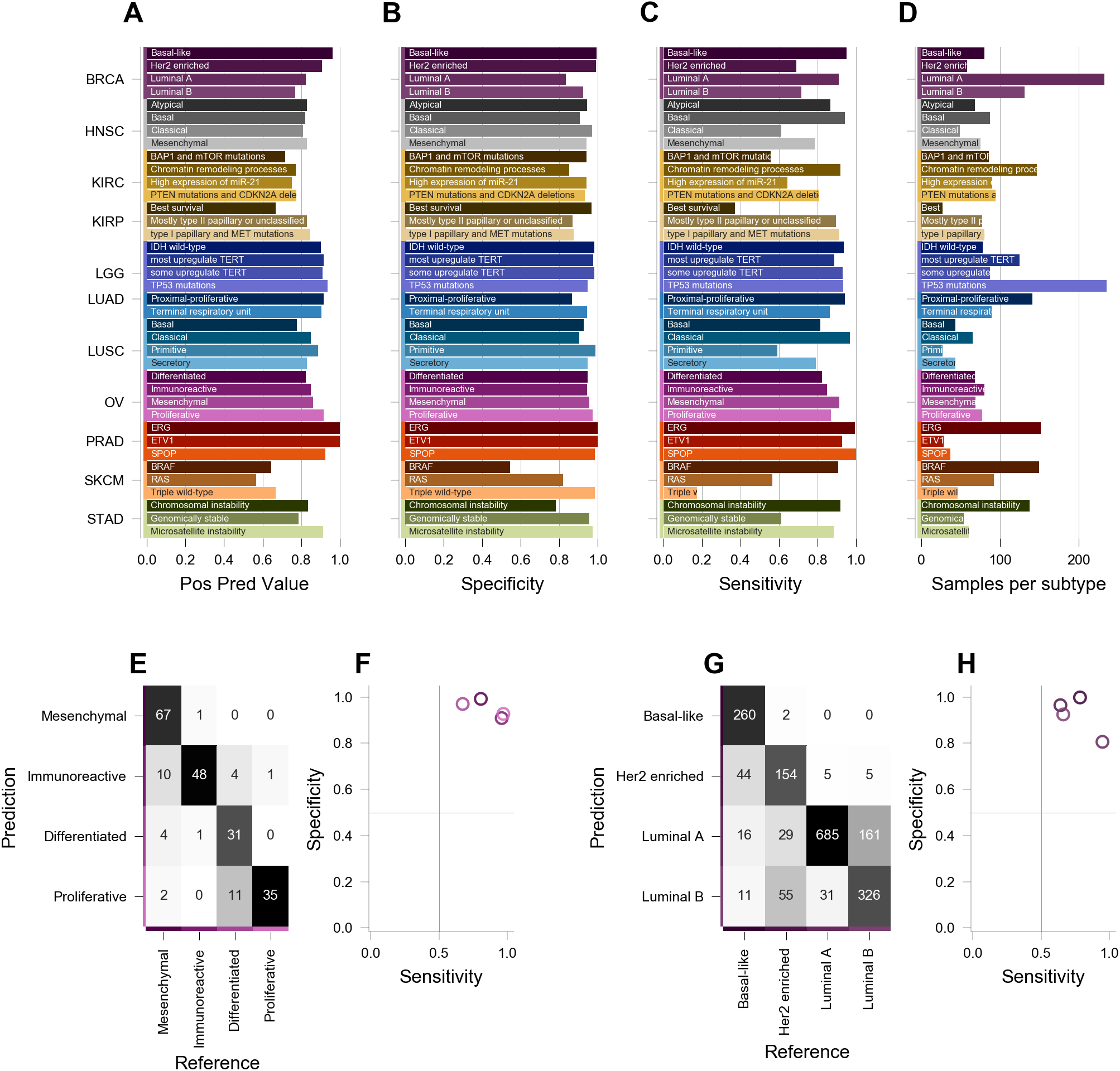
Cross- and external validation of molecular subtype predictors. A predictor of molecular subtypes was constructed for each of 11 primary cancer types, spanning 38 molecular subtypes. (A) Per-class positive predictive value, (B) specificity, and (C) sensitivity of molecular subtype classifications evaluated through cross-validation (Figure 1). (D) Number of training samples for each molecular subtype. To further validate these subtype predictors, breast (E) and ovarian (F) subtype predictors were used to predict the respective molecular subtypes in two external datasets (GSE9899 and EGAS00000000083, respectively). (G-H) Sensitivity and specificity for each classification in (E) and (F), respectively.

#### 3.2.1 Subtype predictors are accurate on external data of different platforms

To further validate the cancer subtype predictors, we classified samples from two external datasets: ovarian cancer [15] and breast cancer [38] annotated with molecular subtypes, with 215 and 1,784 samples respectively. The ovarian cancer subtype predictor (Figure 7E,F) attained a median specificity of 94.9% (the best performance is for mesenchymal: 99.2%) and a median sensitivity of 88.4% (the best performance is proliferative: 97.2%). The breast cancer subtype predictor, shown in Figure 7G,H, presented a median specificity of 95% (the best performance is for basal-like: 99.9%) and a median sensitivity of 72.4% (the best performance is for luminal-A subtype: 95%). Notably, the basal-like molecular subtype of breast cancer is a particularly aggressive subtype and patients relapse rapidly. The accurate classification of this subtype is important for precise treatment—recently, a diabetes drug has been shown as a potential therapy for basal-like breast cancer patients [65]. The two external validation datasets from 1,999 patients with breast or ovarian cancers demonstrate that our subtype predictors can distinguish clinically favorable subtypes from those associated with poor prognosis.

Together with the identification of primary origin for metastatic tumors, followed by subtype classification together allows for informed therapeutic decision making for clinicians, thereby improving treatment outcomes for CUP patients.

## 4. DISCUSSION

It is widely appreciated that cancer is a disease at the scale of the entire genome, but it remains difficult to effectively translate this complexity into clinical utility. Two important pieces of information that are relevant in clinical care and translational research are knowledge of tissue of origin, or CUP, and subtype of the cancer. Identifying tissue of origin of CUPs and molecular subtype is critical for personalized medicine, where the treatment is tailored to the molecular profile of individual tumor [66]. Because of the lack of primary site information, CUP patients receive palliative chemotherapy that lacks the precision of modern targeted cancer medicine and results in no clear benefit in survival [67, 68]. Various approved targeted therapies for cancer by the Food and Drug Administration (FDA) include signal transduction inhibitors, gene expression modulators, apoptosis inducers, angiogenesis inhibitors, immunotherapies. The targeted therapies have been approved for the treatment of over 28 types of cancer. As CUP may retain molecular signatures of its primary site, CUP patients with primary or metastatic tumor might benefit from established therapeutic regimens appropriate for cancers of that tissue; therefore, identifying the primary site is key to choosing effective therapeutic options.

Another major challenge in clinical cancer research is accurately classifying cancers into appropriate homogeneous subtypes to improve prognosis and treatment [5]. Analyses of The Cancer Genome Atlas (TCGA) and the International Cancer Genome Consortium (ICGC) have established that a cancer at any primary site can be further classified into molecular subtypes with potentially distinct clinical outcome and therapeutic options [21–28, 62]. For example, the EBV-positive subtype of gastric cancer is associated with overexpression of JAK2, PD-L1 and PD-L2 genes, suggesting that PD-L1/2 antagonists and JAK2 inhibitors are potential therapeutic options for these tumors [64]. Similar clinical and therapeutic relevance of subtype information has been demonstrated in multiple cancers [21, 23-25, 27, 28]. Increasingly, tumor molecular subtype is being considered as an eligibility criterion for the entry into clinical trials [29]. However, for many cancers, the molecular subtype information is not available for use in clinical practice because of the difficulty in identifying the subtype of a given tumor [30, 31]. Besides being an important factor in clinical decision making, the knowledge of the molecular subtype will be helpful in translational research. For example, cancer avatar trials use PDX models to test panels of drugs to determine the best regime for personalized human therapy. Knowledge of subtype of the tumor may narrow down the choice of treatment regimens to test which increases efficiency of the cancer avatar trials.

We developed predictors of high sensitivity and specificity for classification of primary site of origin of 33 cancers and molecular subtyping of 11 cancers using gene expression data from the TCGA. We show, for the first time to our knowledge, our pan-cancer classifiers can predict multiple cancers’ primary site of origin from metastatic samples. Compared to the other predictors based on somatic mutations [69], our predictors are not limited to only cancer types with high mutation burden and has much greater potential for clinical diagnosis and therapeutic design. Further, the predictors designed based on the TCGA RNA-seq data generalize to different cohorts of primary and metastatic samples profiled using RNA-seq and microarray sequencing technologies. Such external validation qualifies our predictors to be robust across technology platforms, batches and sample processing protocols. A combination of primary tissue of origin and subtype classification from metastases will serve as important tools for clinicians in effectively treating CUPs.

The classification tools discriminated all cancers from each other well, except among the gastro-intestinal cancers. However, the classification by cancer group could be achieved with very high sensitivity and specificity. Different cancers among gastrointestinal cancers are less distinguishable due to their anatomic proximity and molecular similarity, as shown in Figure 3. To circumvent this problem, in our future work, we will adopt a hierarchical classification of tumors: (1) granular classification by organ system (2) finer classification by cancer type in each organ system. In addition, we can include features from copy number, mutation and methylation data to augment our feature set for both accuracy as well as robustness for technology and batch variations. Molecular subtyping can also benefit from adding features from heterogeneous data as several subtypes were identified to exhibit genomic features that span whole spectrum of omics data. For example, the CIN subtype in gastric cancer is known to exhibit large structural variations which may not be captured accurately by expression data. Thus, our future work will encompass comprehensive multi-omic data to identify tissue of origin and molecular subtyping.

Though the overall performance of the predictors designed using DLDA, SVM and KNN is not as good as Random Forest on this data, their performance is on par with or better than Random Forest based predictors on certain tumor type and subtype classification. However, the classification predicted by Random Forest predictors is easier to interpret.

As we continue to enhance the predictor, we do recognize that the clinical utility of the predictors is dependent on their ability to classify FFPE (formalin-fixed paraffin-embedded) samples, which is the standard specimen type used for molecular profiling of cancers in clinical diagnostics in addition to fresh-frozen samples. We have previously shown that the predictors designed to work on microarray data can also work with FFPE samples if they are profiled using nanoString arrays [70]. Therefore, it is feasible to generalize our predictors to work on FFPE samples for clinical applications. In summary, we have demonstrated the utility of gene expression profiles to solve the important clinical challenge of identifying the primary site of origin and the molecular subtype of cancers based on machine learning algorithms. These predictors will be made available as open source software, freely available for academic non-commercial use.

In an effort to make these tools available to as wide an audience as possible, we offer our models and results in two publicly available forms: a web-based portal and a software package which can be used to apply these tools to other datasets and to reproduce the results presented here. The web-based portal provides interactive visualizations showing the expression profiles of the TCGA cancer samples and the classification results of our predictors. These visualizations allow for the exploration of relationships between cancer types in the context of pan-cancer expression profiles.

## SUPPORTING INFORMATION

Table S1. Contingency tables and performance metrics for all primary site predictors

Table S2. Cross-validation performance metrics of subtype predictors

Table S3. External validation performance metrics of subtype predictors

## ACKNOWLEDGEMENTS

Research reported in this publication was partially supported by the National Cancer Institute of the National Institutes of Health under Award Number P30CA034196. The content is solely the responsibility of the authors and does not necessarily represent the official views of the National Institutes of Health. This study makes use of data generated by the Molecular Taxonomy of Breast Cancer International Consortium. Funding for the project was provided by Cancer Research UK and the British Columbia Cancer Agency Branch.

## AUTHOR CONTRIBUTIONS

R.K.M.K. and J.G. designed research. S.L., R.K.M.K. and J.G. guided the project. C.A.P. and J.G. performed data acquisition. W.F.F. and J.G. wrote software and performed analysis. H.R. provided input on clinical oncology and contributed to the interpretation of the results. S.N. provided computational support. W.F.F., S.N., C.A.P., H.R., S.L., R.K.M.K., J.G. wrote the manuscript.

